# Neural Representations of Food-Related Attributes in the Human Orbitofrontal Cortex During Choice Deliberation in Anorexia Nervosa

**DOI:** 10.1101/2021.05.05.441818

**Authors:** Alice M. Xue, Karin Foerde, B. Timothy Walsh, Joanna E. Steinglass, Daphna Shohamy, Akram Bakkour

## Abstract

Decisions about what to eat recruit the orbitofrontal cortex (OFC) and involve the evaluation of food-related attributes, such as taste and health. These attributes are utilized differently by healthy individuals and patients with disordered eating behavior, but it is unclear whether these attributes are decodable from activity in the OFC in both groups and whether neural representations of these attributes are differentially related to decisions about food. We used fMRI combined with behavioral tasks to investigate the representation of taste and health attributes in the human OFC and the role of these representations in food choices in healthy individuals and patients with anorexia nervosa (AN). We found that subjective ratings of tastiness and healthiness could be decoded from patterns of activity in the OFC in both groups. However, health-related patterns of activity in the OFC were more related to the magnitude of choice preferences among patients with AN than healthy individuals. These findings suggest that maladaptive decision-making in AN is associated with more consideration of health information represented by the OFC during deliberation about what to eat.

**Significance Statement:** An open question about the orbitofrontal cortex (OFC) is whether it supports the evaluation of food-related attributes during deliberation about what to eat. We found that healthiness and tastiness information were decodable from patterns of neural activity in the OFC in both patients with anorexia nervosa (AN) and healthy controls. Critically, neural representations of health were more strongly related to choices in patients with AN, suggesting that maladaptive over-consideration of healthiness during deliberation about what to eat is related to activity in the OFC. More broadly, these results show that activity in the human OFC is associated with the evaluation of relevant attributes during value-based decision-making. These findings may also guide future research into the development of treatments for AN.

## 1. Introduction

Deciding what to eat involves the evaluation of multiple types of information and consideration of subsequent consequences and outcomes. Previous studies have shown that the orbitofrontal cortex (OFC) plays a central role in representing the subjective value of individual foods and food choice (Padoa-Schioppa and Assad, 2006; Plassmann et al., 2007; Clithero and Rangel, 2014; Suzuki et al., 2017; Ballesta et al., 2020). Other studies have demonstrated that evaluations of tastiness and healthiness—two food attributes that tend to be unrelated among healthy individuals—interact to determine food choices (Hare et al., 2009, 2011; Maier et al., 2015; Lloyd et al., 2020).

The OFC has been further implicated in the integration of basic food-related attributes during the computation of the subjective value placed on foods (Suzuki et al., 2017), and it is often assumed that these computations also take place during decision-making. But how are these attributes represented at the neural level and how do they contribute to deliberation about what to eat? One approach to addressing these open questions is to compare neural activity and choices in populations that differ in the extent to which they rely on taste versus health attributes when making food-related decisions. Individuals with anorexia nervosa (AN) are well-known to adhere to a low-fat, low-calorie diet even to the point of starvation (Arcelus et al., 2011; Walsh, 2011). Given this well-characterized behavioral profile, examination of the neural representations of taste and health attributes and their link to behavior in AN may offer new insights into the mechanisms that perpetuate this devastating illness. This approach can also facilitate understanding of how food attributes are represented and related to choices, more generally. In the current study, we use multivariate analysis methods to better understand the representations of tastiness and healthiness information in the OFC and how these representations contribute to food choice. Here, we use “taste” to denote subjective ratings of how tasty different foods are and “health” to denote subjective ratings of how healthy different foods are.

Evaluations of taste and health attributes play different roles during food choices among individuals with AN as compared to healthy individuals (Foerde et al., 2015, 2018, 2020; Steinglass et al., 2015, 2016; Uniacke et al., 2020). In an fMRI study, overall levels of activity assessed in univariate analyses of taste and health attribute ratings were differentially associated with choices across individuals, with choice-related ventromedial prefrontal cortex (vmPFC) activity correlated with tastiness-related activity among healthy controls (HC) and healthiness-related activity among patients with AN (Foerde et al., 2015). These findings hint at the possibility that neural representations of taste and health attribute information differentially guide choices in individuals with AN and healthy individuals. Multivariate pattern analysis— which has greater sensitivity than univariate analyses in the detection of mental representations (Norman et al., 2006)—may provide deeper insights into differences between patients with AN and healthy controls (Frank et al., 2016).

We conducted secondary analyses of neuroimaging data from Foerde et al. 2015 using multivariate pattern analyses. In Foerde et al. 2015, participants rated the tastiness and healthiness of a range of different foods and made food choices during fMRI scanning. The goal of the secondary analysis was to more directly test whether taste and health attributes are represented in patterns of brain activity within the OFC. Furthermore, the behavioral relevance of such activity was tested by linking it to individuals’ choices. To do so, we first assessed whether taste and health attribute information could be decoded using multivariate pattern analyses in the OFC during taste and health ratings in both HC and AN (within-task classification). Next, we applied this decoding of taste and health attributes to a subsequent choice phase (cross-task classification) in order to test whether evidence of tastiness- and healthiness-related representations during choices was related to the actual choices made.

## 2. Methods

### 2.1. Participants

Twenty-one hospitalized women with AN and 21 healthy control women (HC) completed this study. In the analyses described below, all HC participants were included. One individual with AN was missing a structural image and excluded from analyses because functional registration could not be performed. This resulted in a final sample of 41 participants.

Participants were right-handed, between the ages of 16 and 39 years old, taking no psychotropic medications, not pregnant, with no history of significant neurological illness, and no contraindication to MRI. HC were normal weight women (BMI between 18 kg/m^2^ and 25 kg/m^2^) and were excluded from participation if they were taking psychotropic medications, had any history of psychiatric illness, or were currently dieting. All participants provided written informed consent and the New York State Psychiatric Institute Institutional Review Board approved the study.

Eating disorder diagnoses were made via the Eating Disorder Examination (EDE) (Fairburn and Terence Wilson, 1993) and co-occurring diagnoses were assessed via the Structured Clinical Interview (SCID) for Diagnostic and Statistical Manual of Mental Disorders, 4th Edition (DSM-IV) (Spitzer et al., 1987). Ten patients met the DSM-5 (American Psychiatric Association, 2013) criteria for the restricting subtype of AN and 11 patients met the criteria for the binge-eating/purging subtype of AN. For participants with AN, study procedures occurred the day after hospital admission. Treatment at NYSPI is provided at no cost for those interested in and eligible for participation. HC received $125 as compensation for their time.

### 2.2. Behavioral Task Procedures

Pre-scan intake was standardized and controlled, as follows: at 12pm, participants were served a research lunch consisting of ~550 kcal (turkey sandwich, Nutrigrain bar, 8 ounces of water). In between lunch and scanning at 2pm, participants were instructed not to eat or drink anything with the exception of water.

Participants completed three tasks in the scanner: taste rating, health rating, and food choice. The order of the taste and health rating tasks was counterbalanced and randomized across participants. Food choices always followed the two rating tasks. Behavioral task procedures are described in detail in Foerde et al., 2015.

#### 2.2.1. Stimuli

Seventy-six food items were presented in each task (**Fig. 1**). Half of the food items were low fat (<30% of total calories from fat, as determined by our staff research nutritionist) and half of the food items were high fat. In each task, the food items were presented on white plates against a black background in high-resolution color photographs. These stimuli are included in the Food Folio by Columbia Center for Eating Disorders stimulus set (Lloyd and Foerde, 2020; Lloyd et al., 2020; Schebendach et al., 2020). The order of stimulus presentation was randomized in each task. A rating scale was shown below the food item on each trial.

**Figure 1.**
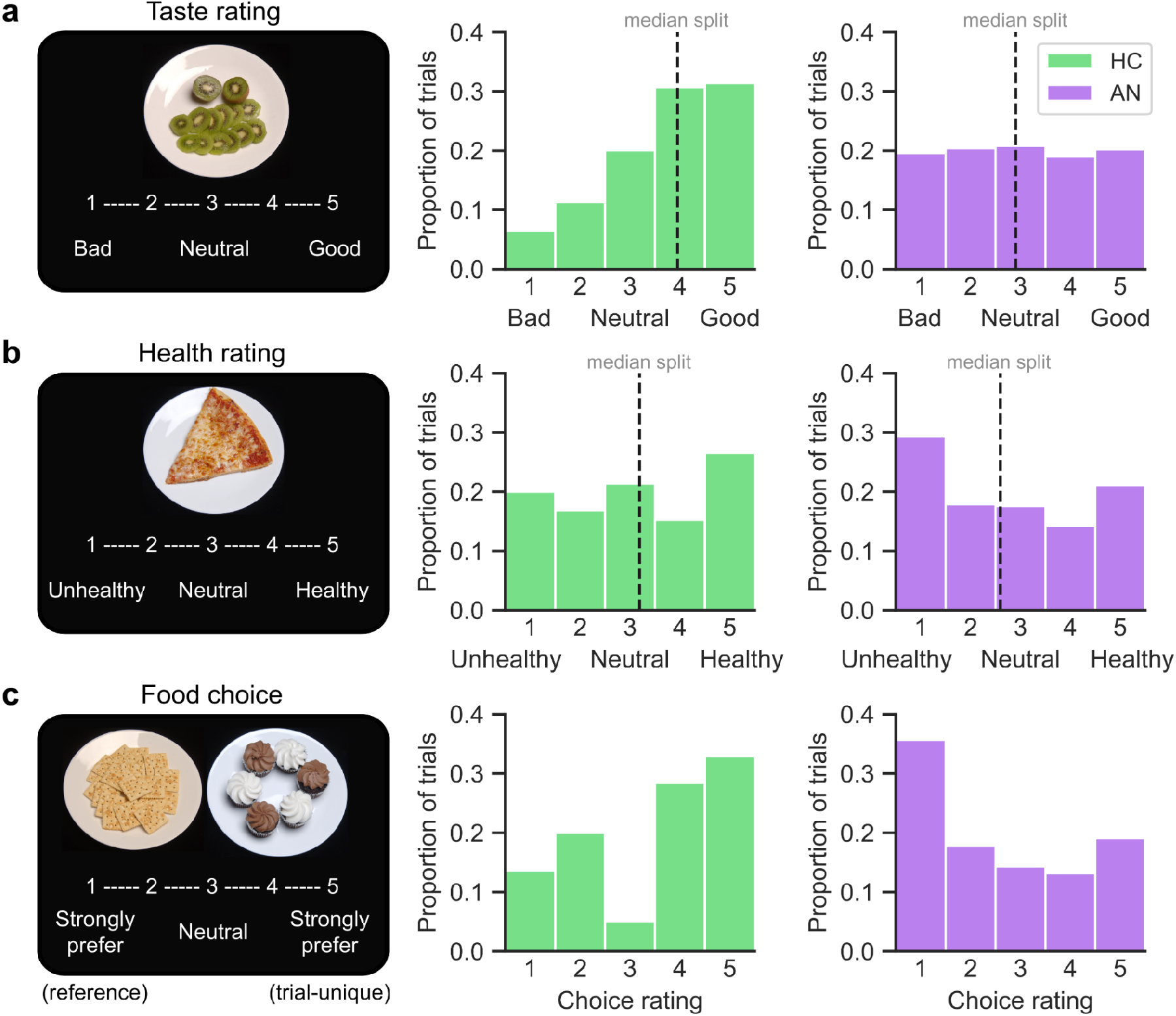
Task design and behavioral results. During taste and health ratings (a and b), participants viewed and rated 76 foods on a Likert scale from 1 to 5. The order of the taste and health rating tasks was counterbalanced across participants. (a) Taste rating distributions are shown for all HC participants in green and all AN participants in purple. Median splits were performed on taste ratings for each participant. The dashed black lines indicate the group-level median across each group of participants (HC=3.95±0.59, AN=2.90±0.83). For the purposes of multivariate pattern analysis, each food was assigned a “low” or “high” taste label according to participant-specific median splits. (b) Health rating distributions are shown for all HC participants in green and all AN participants in purple. Median splits were performed on health ratings for each participant. The dashed black lines indicate the group-level median across each group of participants (HC=3.19±0.60, AN=2.60±0.66). Each food was assigned a “low” or “high” health label according to participant-specific median splits. (c) The rating tasks were followed by a food choice task in which participants were asked to choose between a reference food on the left (rated neutral in taste and health) and a trial-unique food on the right. The reference food was the same on every trial. Participants rated their choice preference on a Likert scale from 1 to 5. The distribution of choice ratings is shown on the right for HC (in green) and AN (in purple).

#### 2.2.2. Taste Rating

In the taste rating task (**Fig. 1a**), participants were asked to rate the tastiness of 76 food items on a five-point Likert scale from “bad” to “neutral” to “good” or “good” to “neutral” to “bad” (the direction of the rating scale was counterbalanced and randomized across participants). They were instructed to rate the food items only on taste.

#### 2.2.3. Health Rating

In the health rating task (**Fig. 1b**), participants were asked to rate the healthiness of 76 food items on a five-point Likert scale from “unhealthy” to “neutral” to “healthy” or “healthy” to “neutral” to “unhealthy” (the direction of the rating scale was counterbalanced and randomized across participants).

#### 2.2.4. Food Choice

The food choice task was completed after the taste and health rating tasks (**Fig. 1c**). For each participant, a reference food item that had been rated by that participant as neutral in taste and health in the rating tasks was selected at random by a computer program. If no food items were rated as being neutral in taste and health, an item that was neutral on health and positive on taste was selected to minimize biasing choices based on taste value. For 20 HC and 18 AN, the reference item was rated by participants as neutral in taste and health. For 1 HC and 1 AN, the reference item was neutral on health and rated 1 step toward “good” on taste. For 1 AN, the reference item was neutral on health and rated 1 step toward “bad” on taste.

During the food choice task, participants were presented the reference food and a trial-unique food on 76 trials. The reference food was always presented on the left side of the screen and was the same on every trial. The trial-unique food was always presented on the right. Participants were instructed to choose the food they would like to eat and indicated their preference on each trial using a Likert scale with “strongly prefer” anchoring each end of the scale. The side-by-side presentation of the foods ensured that participants were aware their choices were relative to the reference food.

To incentivize participants to make choices according to their preferences, participants were told that they would receive a snack-sized portion of one of their chosen foods, selected at random, after the task. Participants were served a snack-sized portion of one of their chosen foods at 3 p.m., observed by staff.

### 2.3. fMRI Acquisition

Neuroimaging was conducted at Columbia University’s Program for Imaging and Cognitive Sciences on a 3.0T Phillips MRI system with a SENSE head coil. Functional data were acquired using a gradient echo T2*-weighted echoplanar (EPI) sequence with blood oxygenation level-dependent (BOLD) contrast (TR=2,000ms, TE=19ms, flip angle=77, 3×3×3mm voxel size; 46 contiguous axial slices). To allow for magnetic field equilibration during each functional scanning run, four volumes were discarded before the first trial. Structural images were acquired using a high-resolution T1-weighted magnetization prepared rapid gradient echo (MPRAGE) pulse sequence.

### 2.4. Imaging Data Preprocessing

Preprocessing of the raw fMRI data was performed using fMRIPrep 1.4.0 (Esteban et al., 2018a, 2018b), which is based on Nipype 1.2.0 (Gorgolewski et al., 2011).

#### 2.4.1. Anatomical Data Preprocessing

The T1-weighted (T1w) image was corrected for intensity non-uniformity (INU) with N4BiasFieldCorrection (Tustison et al., 2010), distributed with ANTs 2.2.0 (Avants et al., 2008), and used as T1w-reference throughout the workflow. The T1w-reference was then skull-stripped with a Nipype implementation of the antsBrainExtraction.sh workflow (from ANTs), using OASIS30ANTs as target template. Volume-based spatial normalization to one standard space (MNI152NLin2009cAsym) was performed through nonlinear registration with antsRegistration (ANTs 2.2.0), using brain-extracted versions of both T1w reference and the T1w template. The following template was selected for spatial normalization: ICBM 152 Nonlinear Asymmetrical template version 2009c (Fonov et al., 2009).

#### 2.4.2. Functional Data Preprocessing

For each of the 3 BOLD runs per subject (across all tasks), the following preprocessing was performed. First, a reference volume and its skull-stripped version were generated using a custom methodology of fMRIPrep. The BOLD reference was then co-registered to the T1w reference using bbregister (FreeSurfer) which implements boundary-based registration (Greve and Fischl, 2009). Co-registration was configured with nine degrees of freedom to account for distortions remaining in the BOLD reference. Head-motion parameters with respect to the BOLD reference (transformation matrices, and six corresponding rotation and translation parameters) are estimated before any spatiotemporal filtering using mcflirt (FSL 5.0.9, (Jenkinson et al., 2002)). The BOLD time-series (including slice-timing correction when applied) were resampled onto their original, native space by applying a single, composite transform to correct for headmotion and susceptibility distortions. These resampled BOLD time-series will be referred to as preprocessed BOLD in original space, or just preprocessed BOLD. The BOLD time-series were resampled into standard space, generating a preprocessed BOLD run in ‘MNI152NLin2009cAsym’ space. First, a reference volume and its skull-stripped version were generated using a custom methodology of fMRIPrep. Several confounding time-series were calculated based on the preprocessed BOLD: framewise displacement (FD), DVARS and three region-wise global signals. FD and DVARS are calculated for each functional run, both using their implementations in Nipype (following the definitions by (Power et al., 2014). The headmotion estimates calculated in the correction step were also placed within the corresponding confounds file. All resamplings can be performed with a single interpolation step by composing all the pertinent transformations (i.e. head-motion transform matrices, susceptibility distortion correction when available, and co-registrations to anatomical and output spaces). Gridded (volumetric) resamplings were performed using antsApplyTransforms (ANTs), configured with Lanczos interpolation to minimize the smoothing effects of other kernels (Lanczos, 1964). Many internal operations of fMRIPrep use Nilearn 0.5.2 (Abraham et al., 2014), mostly within the functional processing workflow. For more details of the pipeline, see <u>the section corresponding to workflows in fMRIPrep’s documentation</u>.

### 2.5. ROI Definitions

We anatomically determined regions of interest (ROIs) using the Automated Anatomical Labeling (AAL) Atlas for SPM12 and transformed them from MNI 6th generation space to MNI152NLin2009cAsym space (Tzourio-Mazoyer et al., 2002; Rolls et al., 2015). The lateral orbitofrontal cortex (lOFC) ROI was created by combining the orbital parts of the left and right middle frontal gyrus, superior frontal gyrus, and inferior frontal gyrus (Suzuki et al., 2017). The medial orbitofrontal cortex (mOFC) ROI was created by combining the medial orbital part of the left and right superior frontal gyrus (Suzuki et al., 2017). The orbitofrontal cortex (OFC) ROI was created by combining the lOFC and mOFC ROIs. The V1 ROI was created by combining the left and right calcarine cortex, as defined by the AAL Atlas. ROIs are displayed in **Fig. 3**.

### 2.6. Imaging Data Analysis (Figure 2)

The classification analyses of interest required several steps. 1) Standard GLM analyses were run to generate the patterns of activity on each trial of each task. 2) Behavioral labels were assigned for every trial and used for classification. 3) Multivariate pattern analysis (MVPA) was used to train a classifier by providing it with patterns of activity in regions of interest along with their labels. 4) The trained classifier was fed a new pattern of activity it had not been trained on to predict the label that ought to be assigned to the pattern. 5) The predicted label was verified as a match with the actual label or not. 6) Steps 3 to 5 were repeated multiple times to determine the classifier’s accuracy. 7) Finally, statistical significance of classification accuracy was determined using non-parametric permutation tests.

**Figure 2.**
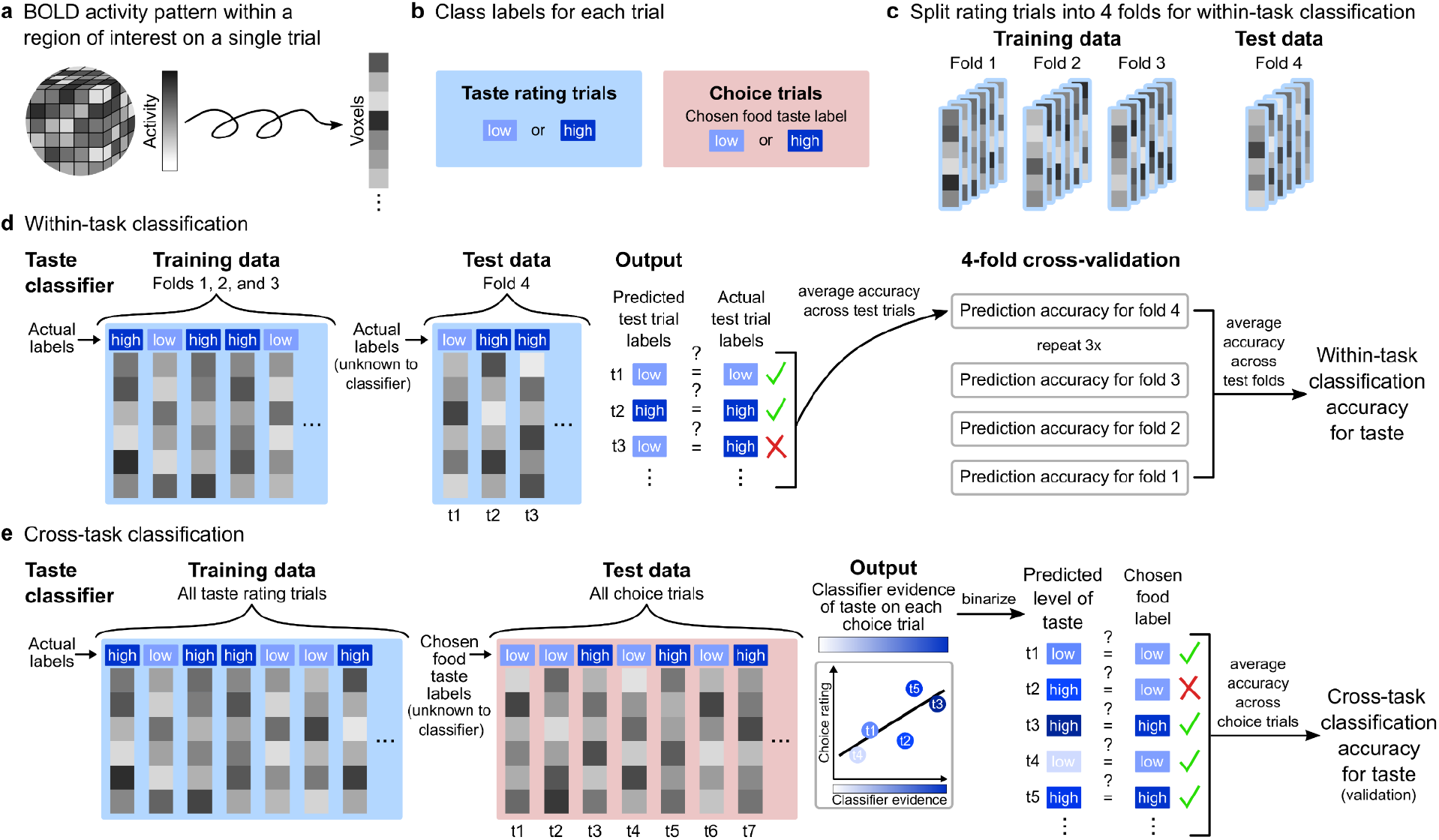
Multivariate pattern analysis approach. (a) Standard GLM analyses were conducted to extract BOLD activity patterns from ROIs from each trial of each task. 3D activity patterns were transformed into vectors of voxel activity, which constituted the features used in subsequent classification analyses. (b) Each trial was assigned a class label. Median splits on taste ratings were conducted for each participant and used to assign each taste rating trial to a high taste class or a low taste class. Each choice trial was then assigned the high/low taste label of the chosen food. The same procedure was followed to assign health class labels to health rating trials and choice trials. (c) For within-task classification of taste, the taste rating trials were split into four partitions. Three folds were used for classifier training and one fold was left out for classifier testing. The same steps were taken for health rating trials. (d) For within-task classification of taste, classifiers were trained on labeled activity patterns from three folds and tested on activity patterns from the left-out fold. The taste classifiers’ predicted high/low taste label for each test trial was compared to actual test trial labels. Classification accuracy for the test fold was defined as mean accuracy across test trials in the corresponding fold. This procedure was repeated three times with a different test fold on each iteration of the crossvalidation procedure. Taste classification accuracy was defined as mean classification accuracy across test folds. Separate taste classifiers were trained and tested for each participant and classification accuracy was averaged across participants in the HC and AN groups. The same procedure was performed for within-task classification of health, except health rating trials and labels were used instead of taste rating trials and labels. (e) The taste classifiers for cross-task classification of taste were trained on labeled activity patterns from all taste rating trials and tested on activity patterns from all choice trials. These classifiers predicted the level of taste evidence in each choice trial’s activity pattern. A linear regression model was run to test the relationship between taste classifier evidence and choice preferences on trials on which the trial-unique item was tasty (taste rating > 3). To validate the cross-task classification approach, the continuous measure of taste classifier evidence was converted to a binary score and compared to the high/low taste label of the chosen food. Cross-task accuracy was defined as mean accuracy in predicting the taste label of the chosen food. Separate taste classifiers were trained and tested for each participant. The same procedures were performed for cross-task classification of health, except health rating trials and labels were used instead of taste rating trials and labels.

#### 2.6.1. GLMs for MVPA Input

We first conducted separate general linear model (GLM) analyses on the preprocessed imaging data for each task to generate input for the multivariate analyses described below. All models were estimated using FSL’s FEAT (Woolrich et al., 2001).

##### 2.6.1.1. GLM Taste

GLM Taste for the taste rating task included three types of regressors: (i) onsets for valid trials (participants responded before the response window ended) were specified by separate regressors, (ii) onsets for timing of the button presses (valid trial onsets + reaction times) were specified by a single regressor, and (iii) onsets for missed trials (participants did not respond within the response window) were specified by a single regressor. On average, HC had 75.2±1.2 valid taste rating trials and AN had 74.0±4.2 valid taste rating trials (out of 76 total). The two groups had a similar number of valid taste trials (t(39)=1.35, p=0.184).

##### 2.6.1.2. GLM Health

GLM Health for the health rating task included three types of regressors: (i) onsets for valid trials (participants responded before the response window ended) were specified by separate regressors, (ii) onsets for timing of the button presses (valid trial onsets + reaction times) were specified by a single regressor, and (iii) onsets for missed trials (participants did not respond within the response window) were specified by a single regressor. On average, HC had 75.6±0.6 valid health rating trials and AN had 74.4±2.0 valid health rating trials (out of 76 total). The number of valid health trials differed significantly between groups (t(39)=2.60, p=0.013).

##### 2.6.1.3. GLM Choice

GLM Choice for the food choice task included three types of regressors: (i) onsets for valid trials (participants responded before the response window ended) were specified by separate regressors, (ii) onsets for timing of the button presses (valid trials onset + reaction times) were specified by a single regressor, and (iii) onsets for missed trials (participants did not respond within the response window) were specified by a single regressor. There were on average 75.4±1.2 valid food choice trials for HC and 74.3±1.9 valid food choice trials for AN (out of 76 total). The number of valid food choice trials differed between groups (t(39)=2.35, p=0.024).

##### 2.6.1.4. GLM Regressors

For all three GLMs, regressors of type (i) were modeled with a boxcar with a duration equal to the trial duration (reaction time), regressor (ii) was modeled with a delta function, and regressor (iii) was modeled with a fixed boxcar with a duration equal to that of the response window (4 seconds). Confound regressors included three translation parameters (in the x, y, and z cardinal planes) and three rotation parameters. As noted in Foerde et al., 2015, quality control analyses indicated that discarding four volumes was insufficient to allow for magnetic field equilibration, so we also included a confound regressor to remove the effects of the first volume by adding a regressor with a 1 for the first volume and 0s elsewhere. No spatial smoothing was applied. All regressors were entered into the first level analysis and all (but the added confound regressors) were convolved with a canonical double-gamma hemodynamic response function. The models were estimated separately for each participant. The parameter estimates for valid trials (regressors of type (i)) were used for subsequent multivariate analyses (**Fig. 2a**).

#### 2.6.2. Multivariate Data Analysis: Within-Task Classification

##### 2.6.2.1. Taste Classification

Decoding analyses were conducted to examine whether taste attribute information was represented in fMRI response patterns in the lOFC and mOFC. A two-class support vector machine (SVM) classifier was trained separately for each participant on patterns of neural activity during taste ratings. This analysis was conducted using PyMVPA with the trade-off parameter (between margin width and number of support vectors) C = 1 (Hanke et al., 2009).

###### 2.6.2.1.1. Definition of Features

The neural activity patterns used as classification samples were raw parameter estimates for the effect of valid rating trials on BOLD (regressors of type (i) from GLM Taste described above). The raw parameter estimate values from voxels within each region of interest (see ROI Definitions above) were the features used to train taste classifiers for each participant (**Fig. 2a and 2d**).

###### 2.6.2.1.2. Definition of Classes

To maximize the number of trials that could be used during training and ensure a balanced number of classification samples in each class, a median split was performed on the taste ratings (**Fig. 2b**). The median taste rating was calculated separately for each participant. Median taste ratings were on average 3.95±0.59 for HC and 2.90±0.83 for AN (**Fig. 1a**). The group-level medians differed significantly between groups (t(39) = 4.71, p < 0.0001). Foods rated below the participant’s median rating were assigned to the “low” taste class. Foods rated above the participant’s median rating were assigned to the “high” taste class. Foods with the median rating value were assigned to the “low” or “high” taste class depending on which assignment minimized the difference in the number of trials between classes. For HC, taste ratings were skewed toward the “good” tasting end of the rating scale, raising concerns that the definitions of high/low taste classes may not have been suitable for participants with skewed taste rating distributions (**Fig. 1a**). For ten out of 21 HC individuals, the “high” taste class only consisted of foods that had the maximum rating of five. For these individuals, classifiers were trained to distinguish “good” tasting foods from “somewhat good,” “neutral,” “somewhat bad,” and “bad” tasting foods. Although “somewhat good” tasting items were placed in the low taste class, cross-validation accuracies were not poorer for HC participants with a median taste rating of five. Instead, a permutation test showed that across ROIs (lOFC and mOFC), cross-validation accuracies for these participants outperformed cross-validation accuracies for participants with lower median taste ratings (p=0.001). Despite many skewed taste rating distributions among HC participants, defining high/low taste classes using a median split produced separable neural activity patterns.

###### 2.6.2.1.3. Cross-Validation Procedure

Classifier training and testing were performed using a four-fold cross-validation procedure (**Fig. 2d**). On each iteration of the cross-validation procedure, the classifiers were trained on three-fourths of taste rating trials. To determine whether the patterns of activity input to the classifiers contained information about taste, we tested whether the trained classifiers could accurately classify each left-out activity pattern from the remaining one-fourth of trials as being high/low in taste. The samples of data used in the left-out partition on each fold were unique and randomly selected. Mean accuracy scores across folds were calculated for each participant and then averaged across participants.

###### 2.6.2.1.4. Determining Statistical Significance

To determine whether taste attribute information was represented in the lOFC and mOFC, the statistical significance of the cross-validation accuracies was tested using permutation tests: the class labels of the trials in the training set were shuffled, four-fold cross-validation was performed, and cross-validation scores were averaged across participants. This procedure was repeated 1000 times to generate a null distribution of mean cross-validation accuracies. For all permutation tests, p-values were the proportion of permuted cross-validation accuracies in the null-distribution greater than the cross-validation accuracies of interest. Mean cross-validation accuracies were considered significant if they were greater than the 95th percentile of the null distribution.

The statistical significance of differences in cross-validation accuracies between groups (HC and AN) and ROIs (lOFC and mOFC) was also tested using permutation tests. A null distribution for group differences was generated by computing group differences 1000 times after shuffling the group labels of cross-validation scores calculated for each participant. The null distribution for ROI differences was generated similarly, but with shuffled region labels instead of shuffled group labels. P-values were calculated as described above.

##### 2.6.2.2. Health Classification

To test whether health attribute information could be decoded from fMRI response patterns in the lOFC and mOFC, the procedures described for taste classification were followed. Any differences in procedure are detailed below.

###### 2.6.2.2.1. Definition of Features

The procedure used to extract neural activity patterns for health classification was identical to the procedure for taste classification, except that raw parameter estimates for the effect of valid rating trials on BOLD were from regressors of type (i) from GLM Health (**Fig. 2a**).

###### 2.6.2.2.2. Definition of Classes

Median health ratings were calculated separately for each participant (group-level median health ratings: HC=3.19±0.60, AN=2.60±0.66, **Fig. 1b**). The group-level medians differed significantly between groups (t(39)=3.05, p=0.004). Foods rated below the participant’s median rating were assigned to the “low” health class. Foods rated above the participant’s median rating were assigned to the “high” health class. Foods with the median rating value were assigned to the “low” or “high” health class depending on which assignment minimized the difference in the number of trials between classes. The suitability of the “low” and “high” class assignments was not assessed here because the health ratings, unlike the taste ratings, were fairly evenly distributed in both groups (**Fig. 1b**).

##### 2.6.2.3. Saturation Classification

Control analyses based on decoding of objective visual information were undertaken to evaluate the classification approach (Suzuki et al., 2017). The analysis steps were identical to those performed for taste and health within-task classification, except for the ROI used. Any deviations in procedures are noted below.

###### 2.6.2.3.1. Definition of Features

The procedure for extracting features were identical to those used for taste and health classification, except these features were extracted from V1.

###### 2.6.2.2.2. Definition of Classes

The saturation of each pixel of each image was extracted using the rgb2hsv function in Matlab. The mean saturation across pixels for each image was used to define “high” and “low” saturation labels according to a median split. This analysis was conducted separately for neural activity during the taste and health ratings.

###### 2.6.2.2.3. Cross-Validation Procedure

The cross-validation procedure for taste and health classification was also performed for saturation decoding from patterns of neural activity during the taste and health ratings.

###### 2.6.2.2.4. Determining Statistical Significance

Permutation tests as described for the taste and health classification analyses were also conducted to assess the statistical significance of saturation decoding. The 95th percentiles were calculated from the resulting taste and health null distributions and p-values were calculated as described above.

##### 2.6.2.4. Exploratory Searchlight Analyses

To examine whether brain regions other than the lOFC and the mOFC contain information about taste and health, we conducted exploratory searchlight analyses. The searchlight analyses were conducted with a searchlight diameter of 5 voxels (i.e. 15mm) and four-fold cross-validation using PyMVPA (Hanke et al., 2009). The resulting searchlight maps were spatially smoothed with a 6mm full width at half maximum (FWHM) Gaussian kernel. To assess the statistical significance of the searchlight maps and to compare the searchlight maps for HC and AN, we used a non-parametric two-sample unpaired t-test against zero and corrected for multiple comparisons using threshold-free cluster enhancement (TFCE) with 5000 permutations. These statistical tests were performed using FSL’s randomise (Winkler et al., 2014). As a control, we also conducted searchlight analyses of saturation, following the same procedure above.

#### 2.6.3. Multivariate Data Analysis: Cross-Task Classification of Taste and Health During the Choice Phase

##### 2.6.3.1. Cross-Task Classification

Once it was determined that taste and health attribute information could be decoded from neural activity patterns in the lOFC and the mOFC, we examined whether neural representations of taste and health attributes were evident in fMRI responses during the choice phase. This classification analysis was conducted using scikit-learn with the regularization parameter C = 1 (Pedregosa et al., 2011).

###### 2.6.3.1.1. Definition of Features

There were no differences between the lOFC and mOFC in the results of the taste and health decoding analyses and subsequent analyses involving the choice task were conducted on the combined OFC ROI (**Fig. 3a and 3b**). Separate taste and health classifiers were trained on all valid taste and health trials for each participant and the classifiers were tested on raw parameter estimates for the effect of valid choice trials on BOLD (regressors of type (i) from GLM Choice described above; **Fig. 2e**).

**Figure 3.**
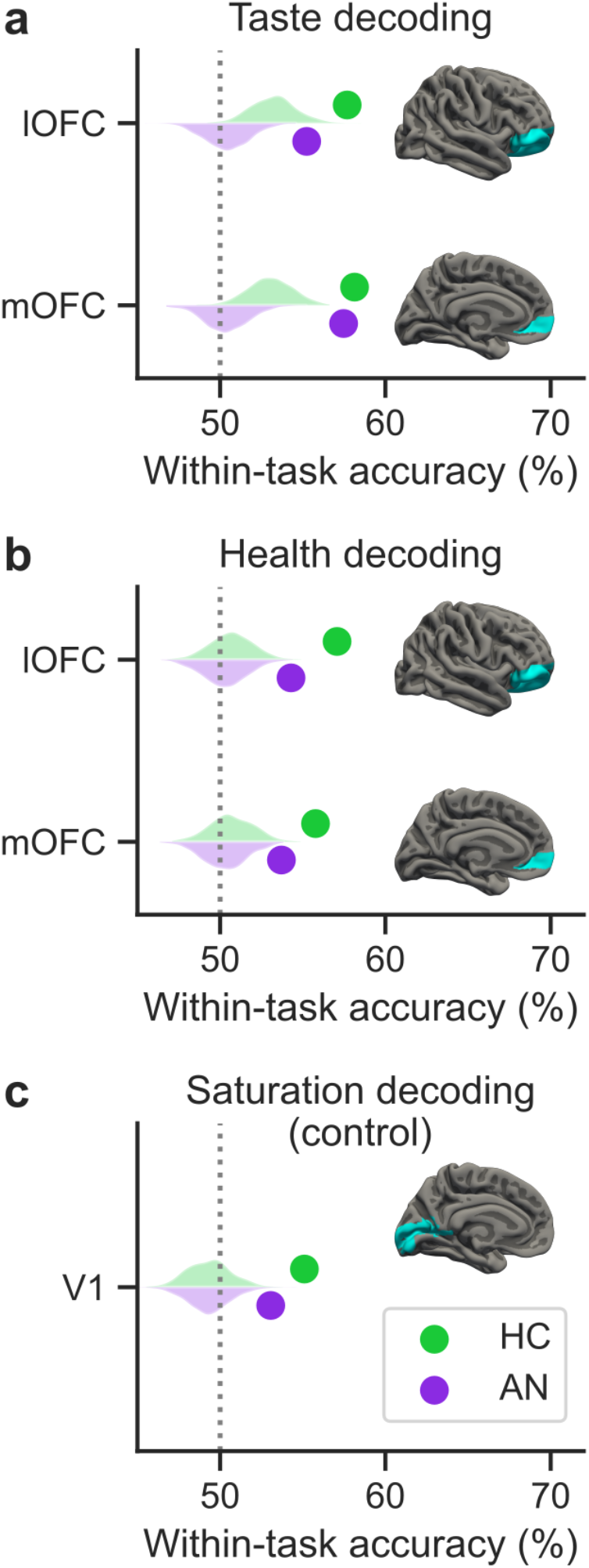
The OFC represents information about tastiness and healthiness. (a) Mean within-task cross-validation accuracy for decoding of tastiness from the lOFC (top) and mOFC (bottom) for HC (in green) and AN (in purple). There were no differences between groups (p=0.21) or subregions (p=0.76). (b) Mean within-task cross-validation accuracy for decoding of healthiness from the lOFC (top) and mOFC (bottom) for HC and AN. Within-task crossvalidation accuracies did not differ between groups (p=0.10) or subregions (p=0.33). (c) Mean cross-validation accuracy for decoding of saturation in V1 from patterns of activity during taste ratings. Similar results were obtained for decoding of saturation from health-related patterns of activity in V1. There were no differences between groups in saturation decoding (from taste rating neural activity: p=0.20; from health rating neural activity: p=0.42). Gray dashed lines indicate chance performance. Violin plots depict the null distributions obtained from permutation tests for each group. The cross-validation accuracies were all greater than the 95th percentiles of the null distributions (all p≤0.004). Anatomically defined regions of interest (lOFC, mOFC, V1) are highlighted in blue in the brain images on the right-hand side of each subplot.

###### 2.6.3.1.2. Cross-Task Classification Output

For each choice task trial, the classifiers output a classifier evidence score between 0 and 1, where a score less than 0.5 indicated evidence of low taste/health and a score greater than 0.5 indicated evidence of high taste/health (**Fig. 2e**). The classifier evidence scores were obtained from the predict_proba function from scikit-learn (Pedregosa et al., 2011).

###### 2.6.3.1.3. Predicting Choice Ratings from Taste and Health Brain Patterns

To examine whether classifier evidence of taste/health information was related to participants’ choices, we ran mixed-effects linear regression models testing the three-way interaction between taste/health classifier evidence scores (continuous), participant group (HC was coded as 0 and AN was coded as 1), and the binarized tastiness/healthiness of the trial-unique item (trials with tasty/healthy trial-unique items (taste/health rating > 3) were coded as 0 and trials with trial-unique items that were not tasty/healthy (taste/health rating ≤ 3) were coded as 1) on choice ratings (1 to 5, with 1 indicating a strong preference for the reference item on the left, 3 indicating no preference, and 5 indicating a strong preference for the trial-unique item on the right). The binarized tastiness and healthiness of the trial-unique item were included in the taste and health models, respectively, to account for the assumption that choice ratings would differ depending on whether the trial-unique item was tastier/healthier than the neutral reference item. More specifically, if participants’ choices were driven by neural evidence of taste, taste classifier evidence should only be positively related to choice ratings when the trial-unique item was tastier than the reference item. Similarly, if choices were driven by neural evidence of health, there should only be a positive relationship between classifier evidence of health and choice ratings when the trial-unique item was healthier than the reference item. In both models, we included a random intercept and random slope for each participant.

###### 2.6.3.1.4. Definition of Classes for Cross-Task Accuracy

During the choice task, two food images were presented simultaneously (**Fig. 1c**), leaving ambiguity about how the images were represented in the neural response. In the choice models relating classifier evidence to behavior, we assumed that the chosen item was more saliently represented than the unchosen item in the neural response and labeled the choice trials with the high/low taste/health of the chosen items (**Fig. 2e**). The alternative possibility, that the trial-specific item (i.e., item on the right) was more saliently represented than the reference item (i.e., item on the left, which was the same on every trial) during each choice trial, was also tested. Here, the choice trials were labeled with the high/low taste/health of the trial-specific items. We also verified that inclusion of the reference item on every trial did not induce novelty preferences over time in the choice phase by testing whether trial number influenced choices for the trial-unique option (no main effect of trial number: odds ratio (O.R.)=0.997, 95% confidence interval (CI)=[0.992, 1.002], p=0.264; no interaction between trial number and group: O.R.=1.00, 95% CI=[0.995, 1.009], p=0.611).

After removing choice trials with a neutral choice rating, on which neither the reference nor the trial-specific item was selected, and trials on which the taste rating of the trial-specific item was not provided, there were 69.9±4.9 choice trials for HC and 60.8±8.1 choice trials for AN (t(39) = 4.39, p < 0.0001). The number of choice trials with health ratings for the trial-specific items and non-neutral choice ratings was 70.3±4.7 for HC and 61.4±8.6 for AN (t(39) = 4.15, p < 0.0002).

###### 2.6.3.1.5. Cross-Task Accuracy Calculation

Classifier evidence scores were converted to binary predictions (scores of less than 0.5 to 0; scores of greater than or equal to 0.5 to 1; **Fig. 2e**) and compared with the high/low taste/health label of the chosen item, as determined by a median split (**Fig. 2b**). To examine whether mean cross-task classification accuracy across participants for each group was significantly above chance performance (50%), we used one-tailed one-sample t-tests. Cross-task accuracies calculated using the ratings of the chosen items and trial-specific items were compared in a mixed-effects linear regression model in R (Bates et al., 2015).

### 2.7. Code Accessibility

Analysis code and outputs are available at https://github.com/alicexue/FCT_MVPA.

## 3. Results

### 3.1. Participant characteristics

The mean age of the HC group was 22.7±3.1 years and the mean age of the AN group was 26.4±6.5 years. Age differed significantly between groups (t(39)=-2.30, p=0.03). The HC group had a mean Body Mass Index (BMI) of 21.5±1.9 and the AN group had a mean BMI of 15.7±2.1. BMI differed significantly between groups (t(39)=9.19, p<0.0001).

### 3.2. Representations of Tastiness and Healthiness in the OFC

#### 3.2.1. Neural Representations in the OFC Reflect Information About Tastiness and Healthiness in Both Healthy Controls and Patients with Anorexia Nervosa

To test whether activity patterns in the OFC reflect high/low taste and health attribute ratings, we trained classifiers to decode high/low taste ratings from brain activity during evaluation of the tastiness of foods and high/low health ratings from brain activity during evaluation of the healthiness of foods. In both groups, taste attribute information could be decoded from the OFC (**Fig. 3a**, all p≤0.001). We found no differences between groups (p=0.21) or subregions of the OFC (p=0.76). Similarly, health attribute information could be decoded from the OFC in both HC and AN (**Fig. 3b**, all p≤0.002), again with no differences between groups (p=0.10) or subregions (p=0.33). Permutation tests indicated that all classification scores were significantly above chance level, and the magnitude of scores was similar to the range of scores reported in a study that employed the same methods (Suzuki et al., 2017). Furthermore, a control analysis decoding objective visual information (saturation) from V1 resulted in mean cross-validation scores that fell in the same range (**Fig. 3c**).

These findings extend prior work showing that the OFC is involved in the evaluation of taste and health attribute information in healthy individuals (Hare et al., 2009, 2011; Londerée and Wagner, 2020) by demonstrating that taste and health attribute ratings could be decoded from neural activity patterns. Additionally, these basic attributes of food could be decoded among individuals with AN, who make very different food decisions.

#### 3.2.2. Neural Representations of Tastiness and Healthiness are Differentially Distributed in Healthy Controls and Patients with Anorexia Nervosa

The representation of tastiness and healthiness throughout the brain in HC and AN was examined in exploratory whole-brain searchlight analyses (**Fig. 4**). Searchlight maps can also be viewed on NeuroVault (https://neurovault.org/collections/MHPZTYJS/). Taste attribute information was decodable from more brain regions among HC compared to AN, and tastiness decoding in HC outperformed tastiness decoding in AN in several regions. The lOFC and mOFC were included among the regions from which tastiness decoding in HC outperformed tastiness decoding in AN. The distribution of above-chance accuracy for healthiness decoding across the brain did not differ significantly between groups. These analyses suggest that outside of the OFC, there are differences between groups in the decodability of tastiness information but not healthiness information.

**Figure 4.**
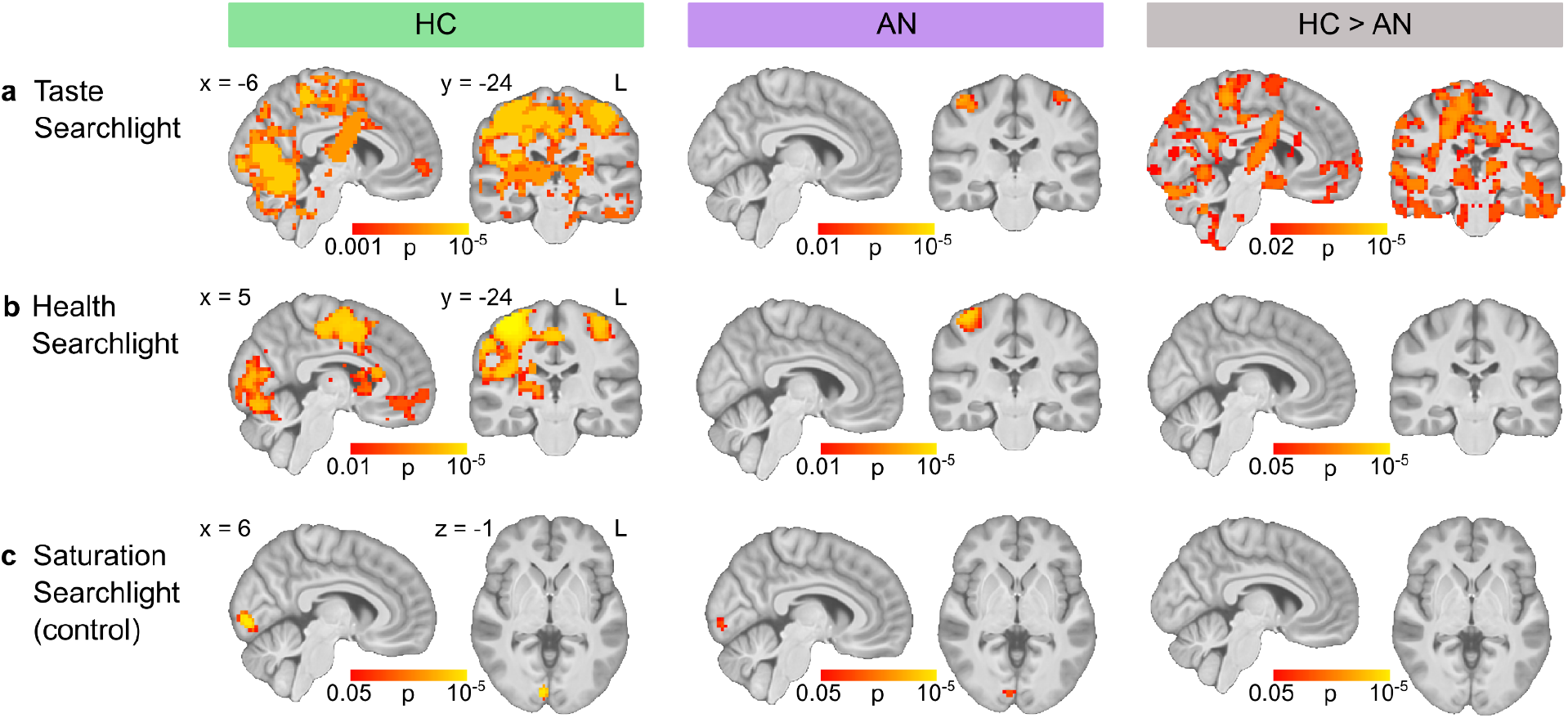
Whole-brain searchlight maps for tastiness and healthiness decoding. Wholebrain searchlight maps for (a) within-task taste attribute decoding, (b) within-task health attribute decoding, and (c) saturation decoding from the taste rating task. Decoding results for HC are displayed below the green column heading. Decoding results for AN are displayed below the purple column heading. Maps depicting where decoding accuracy in HC was greater than decoding accuracy in AN are shown below the brown column heading. Coordinates are reported in MNI152 space. Color bars indicate statistical significance.

### 3.3. Relationship Between Neural Representations of Tastiness/Healthiness and Choice Behavior

#### 3.3.1. Neural Representations of Health Are More Strongly Related to the Magnitude of Choice Preferences in Patients with Anorexia Nervosa Than in Healthy Controls

Participants provided continuous responses in the behavioral choice task indicating how much they preferred to eat the reference item (i.e., item neutral in taste and health, the same on every trial, and presented on the left) or the trial-unique item (i.e., item on the right) (**Fig. 1c**). We sought to examine the extent to which neural evidence of taste/health attribute information in the OFC was related to the magnitude of food choice preferences by entering the continuous choice responses into mixed-effects linear regression models. In these models, we included a binary factor indicating whether the trial-unique item was tasty/healthy (see **2.6.3.1.3**) to account for the assumption that choice ratings would depend on whether the trial-unique item was tastier/healthier than the neutral reference item.

Greater taste classifier evidence in the OFC in HC was not associated with a stronger preference for tasty trial-unique items (no main effect of classifier evidence; **Fig. 5a**; **Table 1**) and the relationship between taste classifier evidence and choice preferences did not differ between groups (no interaction between classifier evidence and group; **Fig. 5a**; **Table 1**).

**Figure 5.**
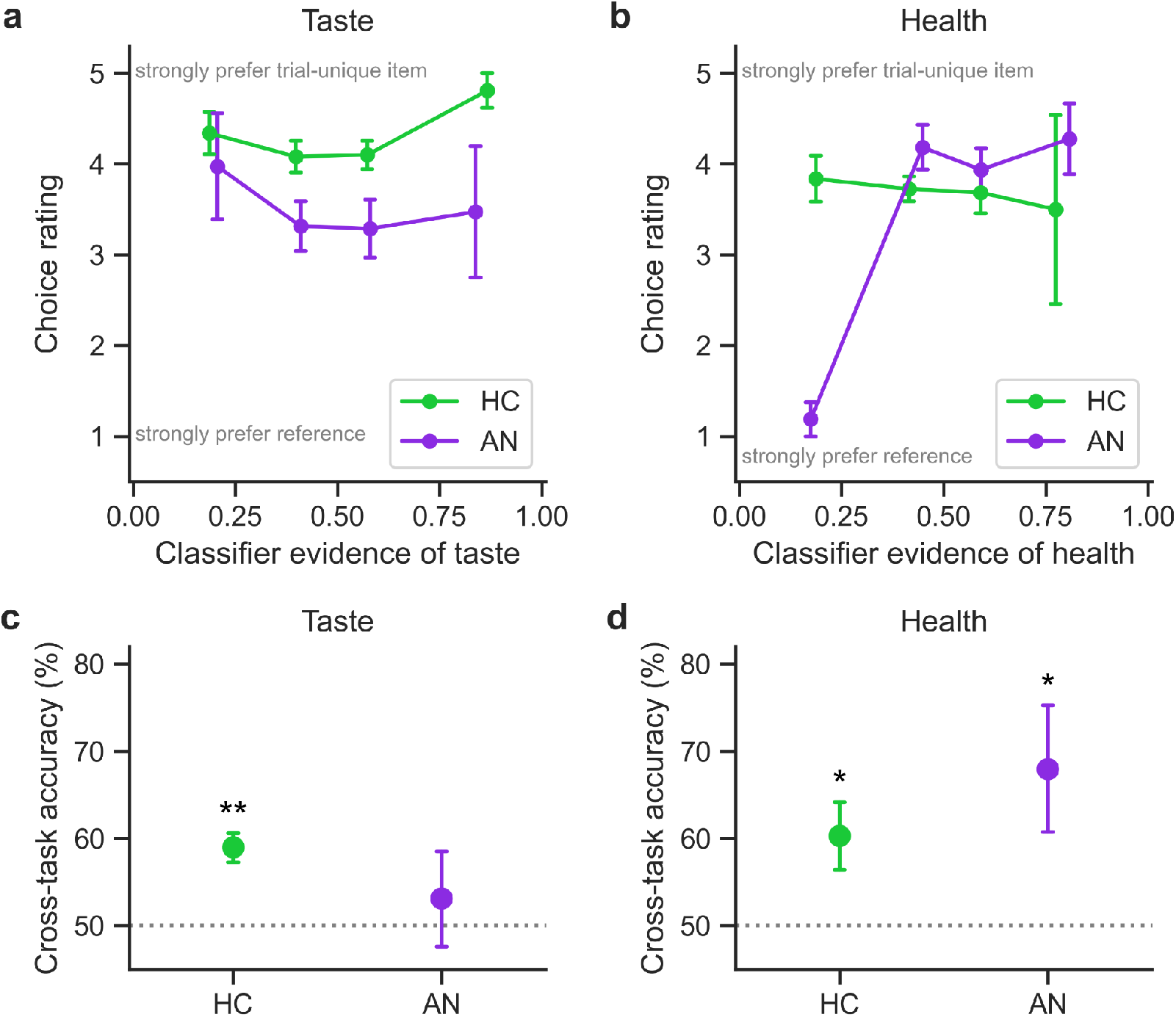
Health classifier evidence in the OFC was related to the magnitude of choice preferences in patients with anorexia nervosa. (a) On trials with tasty trial-unique items (taste rating > 3), taste classifier evidence was not related to HC participants’ preference for the trial-unique option. The relationship between taste classifier evidence and choice preferences did not differ between HC and AN (**Table 1**). (b) On trials with healthy trial-unique items (health rating > 3), health classifier evidence and choice preferences were more strongly related for AN compared to HC (**Table 1**). Plots in (a) and (b) depict mean choice ratings across participants for binned classifier evidence (error bars indicate standard error of the mean). (c) Cross-task accuracy for the taste of the chosen item was significantly above chance for HC (in green; t(20)=5.27, p<0.0001) but not AN (in purple; t(19)=0.53, p=0.302). (d) Cross-task accuracy for the health of the chosen item was significantly above chance for HC and AN (HC: t(20)=2.56, p=0.009; AN: t(19)=2.44, p=0.012). In (c) and (d), error bars indicate standard error of the mean. **p<0.001, *p<0.05.

**Table 1.**
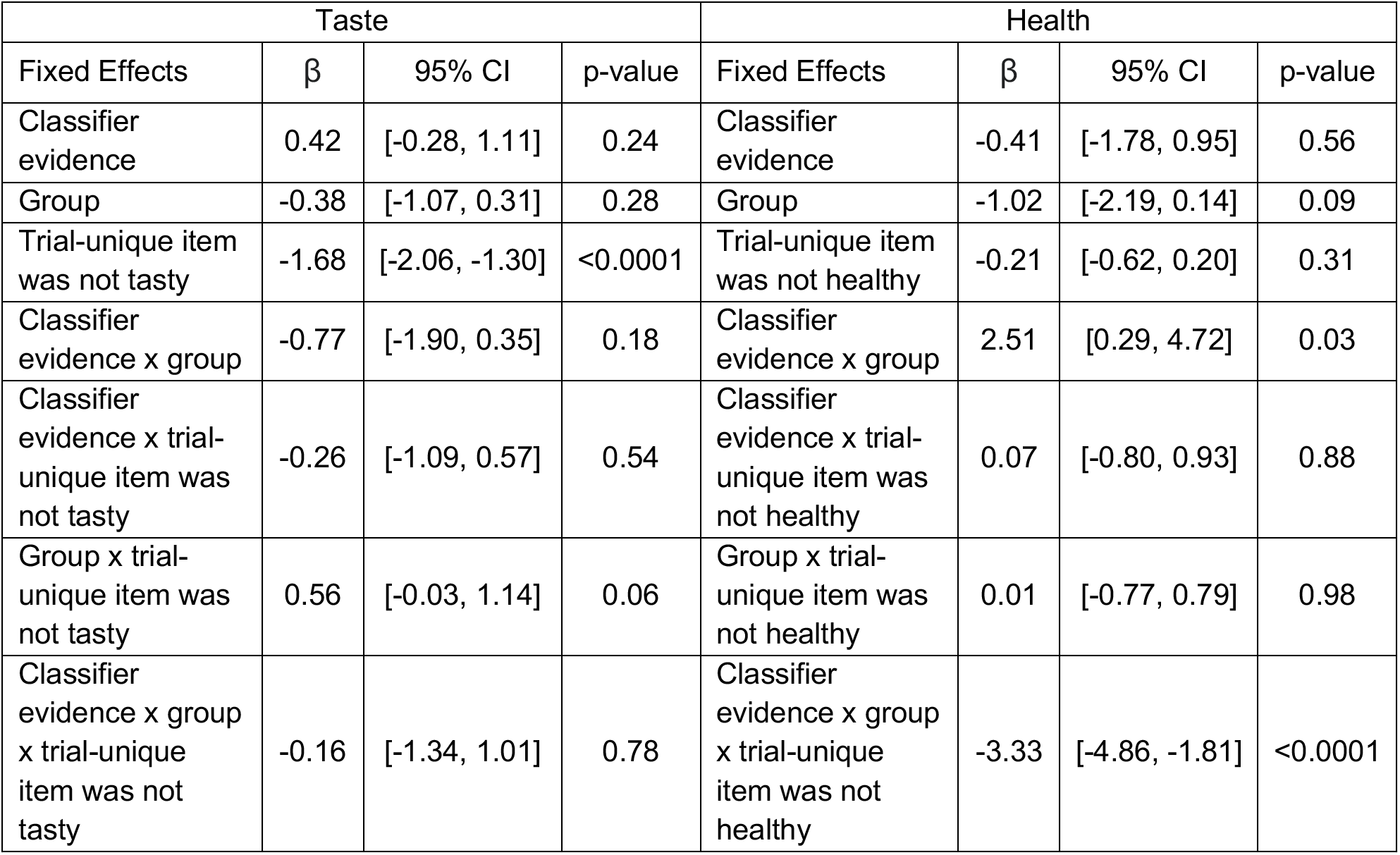
The Effects of Classifier Evidence and Group on the Magnitude of Food Preferences. The relationship between taste/health classifier evidence and choice preferences. For the group variable, HC was coded as 0 and AN was coded as 1. In the taste model, the binarized tastiness of the trial-unique item was coded as follows: trials with tasty trial-unique items (taste rating > 3) were coded as 0 and trials with trial-unique items that were not tasty (taste rating ≤ 3) were coded as 1. In the health model, the binarized healthiness of the trial-unique item was coded as follows: trials with healthy trial-unique items (health rating > 3) were coded as 0 and trials with trial-unique items that were not healthy (health rating ≤ 3) were coded as 1.

There was a stronger positive relationship between health classifier evidence in the OFC and choice preferences for AN compared to HC on trials with healthy trial-unique items (significant interaction between classifier evidence and group; **Fig. 5b**; **Table 1**). Health classifier evidence in the OFC was not related to a stronger preference for healthy trial-unique items in HC (no main effect of classifier evidence; **Fig. 5b**; **Table 1**). These findings provide support for the idea that the OFC plays an important role in the over-consideration of health information during maladaptive decision-making.

#### 3.3.2. Control Analyses to Validate Cross-Task Classification

We sought to establish that taste/health classifier evidence was a meaningful measure to assess during the choice phase, as participants were not instructed to consider tastiness or healthiness when making their choices, and participants viewed two items on each trial of the choice task and only one item on each trial of the rating tasks. Because the cross-task classifiers’ output for each choice trial was a continuous measure of evidence of taste/health, there was ambiguity as to which item this evidence was reflective of. To assess the validity of the cross-task classification approach, we tested whether the level of classifier evidence on each trial was reflective of the label of the chosen food (our assumption) or, alternatively, the label of the trial-unique food, neither of which was not known to the classifiers (**Fig. 2e**).

Cross-task accuracy for the tastiness of the chosen food was defined as the proportion of trials on which the high/low level of taste classifier evidence matched the high/low taste label of the chosen food. Note that the chosen food could have been the reference food (i.e., the same item always presented on the left) or the trial-unique food (i.e., the item on the right). Cross-task classification accuracy for the tastiness of the chosen food was significantly above chance for HC (t(20)=5.27, p<0.0001), but not AN (t(19)=0.53, p=0.302) (**Fig. 5c**). Similarly, cross-task accuracy for the healthiness of the chosen food was defined as the proportion of trials on which the high/low level of health classifier evidence matched the high/low health label of the chosen food. Cross-task classification accuracy for the healthiness of the chosen food was significantly above chance for both HC (t(20)=2.56, p=0.009) and AN (t(19)=2.44, p=0.012) (**Fig. 5d**).

We also tested the alternative possibility that taste/health classifier evidence from patterns of neural activity during choices reflected attributes of the trial-specific items more so than those of the chosen items. Cross-task classification of the tastiness and healthiness of the chosen items outperformed cross-task classification of the tastiness and healthiness of the trial-specific items (main effect of chosen/trial-specific item on cross-task accuracy: β=6.01, 95% confidence interval (CI)=[1.22, 10.79], p=0.015). Because trial-specific items were always presented on the right-hand side of the screen, this result also suggests that the classifiers were not simply decoding leftward or rightward responses. Although the order of the rating scales was counterbalanced across participants, we further assessed the possibility of a left/right response confound by relating classifier evidence to the participants’ choices for the item on the right-hand side of the screen. Classifier evidence of taste did not predict choices for the item on the right side of the screen (no main effect of classifier evidence: odds ratio (O.R.)=2.28, 95% CI=[0.65, 8.03], p=0.20) and this relationship did not differ between groups (no interaction: O.R.=0.20, 95% CI=[0.03, 1.25], p=0.09). Similarly, classifier evidence of health did not predict choices for the item on the right side of the screen (no main effect of classifier evidence: O.R.=0.96, 95% CI=[0.13, 7.10], p=0.96) and this relationship did not differ between groups (no interaction: O.R.=1.25, 95% CI=[0.05, 28.38], p=0.89). Together, these findings suggest that attributes of the chosen food are reflected in the measure of classifier evidence and support the validity of the cross-task classification approach.

## 4. Discussion

The current study examined representations of key food-related attributes—taste and health—in the OFC, and the role of these representations in food choice. Taste and health attribute information were represented in the OFC not only among healthy individuals, but also among patients with AN. The latter routinely make very different and maladaptive food choices as compared with HC (Hadigan et al., 2000; Steinglass et al., 2015; Schebendach et al., 2019). Notably, information about subjective ratings of health in the OFC was a better indicator of the magnitude of choice preferences among individuals with AN than HC. These findings demonstrate that representations of health in the OFC are differentially related to normative and maladaptive decisions about food.

Neuroimaging studies investigating the OFC in AN have focused primarily on structural alterations. Higher gray matter volume in the mOFC among patients with AN relative to HC (Frank et al., 2013; Lavagnino et al., 2018) raises the question of whether informational content in this region differs between groups. Restrictive eating among patients with AN could potentially result from amplified representations of health attribute information or diminished representations of taste attribute information. In our ROI-based analyses, there were no differences between HC and AN in the decodability of either type of information from the OFC. In a whole-brain searchlight analysis, tastiness was significantly more decodable from several brain regions among HC compared to AN, suggesting that this information is not selectively represented in the OFC. These diminished representations of the tastiness of foods in AN extend prior work showing weaker representations of gustatory information in this clinical population (Frank et al., 2016).

Converging evidence suggests that informational content in the OFC differs along the medial-lateral axis. More specifically, different OFC subregions are thought to have distinct roles in supporting value-based decisions, such that the lOFC encodes identity-specific attributes and the mOFC encodes general value (Howard et al., 2015; Suzuki et al., 2017; Vaidya and Fellows, 2020; Howard and Kahnt, 2021). Although these findings may suggest that taste and health attributes are selectively represented in the lOFC, we did not find differences between OFC subregions in decoding accuracy for taste or health attribute information in HC. Among patients with AN, there was a similar pattern of results. Taste and health attributes may have comparable representations in these OFC subregions because they each encompass the values of a wide array of food characteristics (Lloyd et al., 2020). Unlike specific nutritive attributes (Suzuki et al., 2017), or the sweet and savory qualities of foods (Howard et al., 2015), taste and health attributes may be invariably represented in value signals in both the lOFC and the mOFC (Hare et al., 2014; Lloyd et al., 2020). However, our ability to address these questions is limited because the current study was not designed to probe differences in informational content in different subregions of the OFC.

Here, we used multivariate pattern analysis techniques to assess whether the diminished influence of taste attributes and enhanced influence of health attributes on food choices among patients with AN are related to patterns of neural activity in the OFC. Previous univariate analyses of this dataset suggested that taste representations in the vmPFC influence choices in HC while health representations in the same region influence choices in AN (Foerde et al., 2015). Contrary to our expectations, we did not find that neural evidence of taste attribute information in the OFC was related to a stronger preference for tasty items in normative decision-making. This could potentially be explained by the skewed distribution of taste ratings among healthy individuals. Ten out of 21 HC participants had a high-taste class that only included foods with a rating equal to the maximum rating of five. Lower levels of taste classifier evidence for these HC participants could be indicative of representations of tastiness that correspond to a rating of four (somewhat tasty). This reduced sensitivity to tastiness may explain why classifier evidence of taste among HC did not predict the magnitude of their choice preferences. Future studies that employ food stimulus sets with more normally distributed taste ratings among participants may have more success in characterizing the contribution of OFC representations of tastiness to choices in normative decision-making (Lloyd et al., 2020).

For health information, there was a stronger brain-behavior relationship in AN compared to HC. Neural evidence of health attribute information in patterns of activity in the OFC was predictive of the magnitude of choice preferences made by individuals with AN, suggesting that the OFC has a fundamental role in using health information to guide food choices among these patients. This complements behavioral findings that patients with AN—more so than healthy individuals— consider health more strongly in their food-related choices (Foerde et al., 2015, 2020; Steinglass et al., 2015, 2016; Uniacke et al., 2020). These findings point to an important role for the OFC in the over-consideration of health information during maladaptive decision-making in AN and complement previous work showing that the vmPFC is related to the ability to exert executive control and bias the influence of food-related attributes on choice in healthy individuals (Hare et al., 2009; Maier et al., 2015). Studying how health information is learned and encoded by patients with AN may provide additional insights into the neural mechanisms underlying maladaptive decision-making in this disorder. The contribution of hippocampal-based memory systems to the retrieval of knowledge about elemental attributes of foods during food valuation may be an interesting avenue of future research (Barron et al., 2013; Tang et al., 2014; Bakkour et al., 2019).

Whereas the encoding of subjective value in the OFC has been well-established (Levy and Glimcher, 2012), the process by which value-based food choices are made remains disputed. One prevailing theory posits that the OFC supports preference-based decisions by integrating value-predictive attributes into an abstract, common currency for comparison (Wallis, 2007; O’Doherty et al., 2021). Alternatively, relevant attributes may be directly compared in the OFC (Perkins and Rich, 2021). Although the current study was not designed to test whether information about food attributes is integrated or simply compared, we found evidence of taste and health attribute representations in the OFC during choices among healthy individuals. Future studies with tasks specifically designed to address these theories, along with the cross-task multivariate analysis approach taken here, may be able to elucidate (a) whether choice option attributes, like healthiness, are compared during choice deliberation and (b) whether these comparative processes support value construction.

Studies of food choice that have assessed decision-making as a function of tastiness and healthiness (Hare et al., 2009, 2011; Foerde et al., 2015; Maier et al., 2015) generally assume that weighted sums of these attributes are computed to guide decisions. Here, the taste and health rating tasks were used to obtain subjective ratings of these attributes (Hare et al., 2009, 2011; Sullivan et al., 2015; Lloyd et al., 2020; Maier et al., 2020). Although participants were instructed in the taste rating task to only evaluate foods on taste, assessment of how good or bad different foods taste could potentially be thought of as similar to value assignment, during which features other than palatability are considered (Suzuki et al., 2017). Although participants were not asked to report their interpretation of the taste rating task instructions, we follow previous studies in conceiving of the tastiness ratings as measurements of a single component of food value (Hare et al., 2009, 2011; Maier et al., 2020).

Subjective value judgments, unlike judgments of tastiness, are sensitive to personal goals and context. Among healthy individuals, value signals in the vmPFC are responsive to long-term dietary goals and cues to attend to the healthiness of foods (Hare et al., 2009, 2011). The relationship between tastiness and value, however, has been somewhat challenging to characterize in healthy individuals, as previous findings indicate that drawing attention to tastiness does not enhance the influence of this attribute on choices or neural value signals (Hare et al., 2011; Tusche and Hutcherson, 2018). In the study of decision-making among individuals with AN, the distinction between these concepts is particularly important to consider because this clinical population is less influenced by tastiness in their food choices (Foerde et al., 2015; Steinglass et al., 2015; Uniacke et al., 2020). Contrary to our expectations, we did not find differences between groups in the role of tastiness neural evidence in the OFC in choice ratings. The mechanisms by which tastiness exerts less influence on subjective value in AN thus remain a largely unexplored field of inquiry. Future work studying whether tastiness information is more strongly incorporated in OFC value signals in obesity or addiction could provide insights into the role of tastiness evaluation in subjective value computations more generally. This line of research on the role of attribute consideration in value-based decisionmaking may also benefit from investigations of the evolution of value signals and attribute representations in the OFC over the course of choice deliberation (Sullivan et al., 2015; Motoki et al., 2018; Maier et al., 2020).

The present study was not specifically designed to undertake the analyses presented here and some limitations warrant consideration. The distribution of taste ratings among HC was skewed relative to AN, resulting in high/low-taste food labels that did not capture the well-characterized influence of taste on choices among HC that was also previously observed in this sample (Foerde et al., 2015). This did not appear to alter the decoding results presented here, but future studies that seek to employ classification methods could specifically select stimulus sets to address such concerns (Lloyd and Foerde, 2020; Lloyd et al., 2020; Schebendach et al., 2020). It should be noted that the magnitude of decoding accuracies should be interpreted with caution because these measures can be influenced by ROI size, degree of voxel smoothing, and the size of training and test data sets, among other factors (Haynes, 2015). As is the case in most decoding studies, the question of interest concerned the prevalence of specific information in certain ROIs. Non-parametric permutation tests, which provide more valid population-level inference than t-tests, revealed that taste and health attribute information were decoded from lOFC and mOFC significantly above chance (Allefeld et al., 2016).

The results from the present study contribute to understanding the valuation process undertaken during food choices and provide insight into differences in the neural mechanisms that support how information about food is used during decision-making in healthy individuals and maladaptive decision-making in AN. These findings point to the importance and complexity of health information in food choice in AN. Recent advances in real-time fMRI neurofeedback technology or neuromodulation (e.g., rTMS) can perhaps be used in conjunction with multivariate analysis methods as promising avenues for understanding the mechanisms underlying the use of health and taste attributes to guide food choices in individuals with AN (Thut and Pascual-Leone, 2010; Watanabe et al., 2017; Dalton et al., 2020). Ultimately, the aim of this work would be to downgrade the importance of health during food-related decisions among patients with AN. The current findings indicate that food decisions involve balancing different attributes of choice options (like health and taste), and that over-consideration of one attribute over others may cause disruptions in choice behavior and lead to persistent maladaptive behavior.

## Acknowledgements

This work was funded in part by the Global Foundation for Eating Disorders, the US National Institute for Mental Health (R01 MH079397, K23 MH076195, K24 MH113737), the National Science Foundation (grant 1606916), the McKnight Foundation, and the Klarman Family Foundation.

